# Adapting Upright Light Sheet Fluorescence Microscopy for Imaging at Air-Liquid Interface

**DOI:** 10.64898/2026.04.07.716945

**Authors:** Chad M. Hobson, Kenji Izumi, Jesse S. Aaron, Navaneetha Krishnan Bharathan, M. Fernanda Ceriani, William Giang, Juan I. Ispizua, Andrew P. Kowalczyk, Rachel M. Lee, E. Angelo Morales, Owen F. Puls, Ellen Quarles, Micaela Rodriguez-Caron, Sara N. Stahley, Francisco Tassara, Shaohe Wang, Shunze Yao, Takanori Tsuchiya, Teng-Leong Chew

## Abstract

Light sheet fluorescence microscopy (LSFM) is increasingly appreciated as the gold standard for gentle, volumetric imaging with fast acquisition speeds and/or long imaging durations. However, the often-constrained sample space of these microscopes has precluded a specific class of biological specimens from being studied with these tools: those requiring an air-liquid interface (ALI). Here, we present a device for robust imaging at ALI on an upright light sheet microscope with dipping objectives. We demonstrate the system using three relevant use-cases: *ex vivo* embryonic mouse salivary glands, human epidermal equivalent cultures, and *in vivo* adult *Drosophila melanogaster* brains. While the device presented is engineered for one specific light sheet microscope design, it provides a blueprint for easy adaptation to other systems. In doing so, it can potentially spur the use of LSFM for model systems that have so far been unable to take advantage of this powerful technology.

## Introduction

Light sheet fluorescence microscopy (LSFM) provides optical sectioning in 3D biological tissue, comparable to that of confocal microscopy, but at a fraction of the light dosage and increased imaging speeds^1–3^. By restricting the excitation light to a thin plane, detection efficiency is maximized while overall light exposure is minimized. Together, this constitutes a means of generating high-contrast fluorescence images with comparatively low levels of photodamage – an ideal combination for live imaging of dynamic processes. LSFM can be performed in a myriad of configurations as is reviewed previously^3–6^, each with unique pros and cons. Here, we turn our attention to the upright LSFM configuration, such as that of the well-disseminated inverted selective plane illumination microscope (iSPIM)^7^, dual-view iSPIM (diSPIM)^8,9^, and the original implementation of the lattice light sheet microscope (LLSM)^10^. These methods introduce two objective lenses from above the specimen oriented at 90⁰ to each other. Importantly, both excitation and detection lenses are dipped into a bath of imaging medium along with the specimen. Such an approach is advantageous because it simultaneously (i) allows for a minimal detection path and therefore maximal detection efficiency, (ii) can be modified for high-resolution or multi-view imaging, (iii) enables coverslip-based sample mounting, (iv) maintains objective working distance for samples that fit between the objective lenses, and (v) minimizes depth-dependent aberrations. For these reasons, upright LSFM has been used to study embryogenesis in *Caenorhabditis elegans*^11^, trans-endothelial neutrophil migration^12^, and cytoskeletal rearrangement of organelles during mitosis in cultured cells^13^, among many other biological processes. However, the wider application of upright LSFM can be limited due to the physical constraints of the sample space.

One such limiting constraint is that upright light sheet configurations require the specimen to be fully submerged in imaging medium. While this is amenable to many model systems such as zebrafish, cultured cells, organoids, and *Drosophila* embryos, it has also precluded many important biological studies. For example, many epithelial tissues such as skin^14^, eyes^15^, lungs^16^, and airways^17^ must be cultured such that only the basal surface is exposed to medium while the apical surface is in air. Alternatively, certain organ explants require a similar air-liquid interface (ALI) for gas exchange and oxygenation to enable proper development^18–25^. Without proper maintenance of this air-liquid interface, live imaging experiments are then resigned to exceptionally short durations. Moreover, non-aquatic whole animals are incompatible with liquid immersion imaging. Live imaging of such model systems and tissues is therefore restricted to more conventional microscopy systems, thus hindering the progress of biological research into their form and function.

Here, we address this limitation by presenting a novel device for upright LSFM at ALI (LSFM-ALI). We demonstrate the device by live imaging three broad classes of air-liquid interface samples: (i) those predominantly in medium with one side exposed to air, (ii) those predominantly in air with one side exposed to medium, and (iii) developed organisms that require air for respiration. In each case, we present live, multi-color, volumetric, time-series LSFM of specimens that were previously not compatible with such microscopes. The device constitutes an important step forward in bringing the benefits of LSFM to suites of tissues and model systems that extend beyond the canonical few.

## Results

### Air-Liquid Interface Device for Upright Light Sheet Fluorescence Microscopy

Here, we present our design for the LSFM-ALI device (Figure 1), the basis of which is to create a pocket of air within a bath of imaging media. To do so, we use a flat stainless-steel base (silver), upon which a silicone spacer (pink) is sealed. A plastic ring (blue) is placed into the silicone spacer, leaving a central circular opening at which an air-liquid interface can be formed. Depending on the experimental goals, a membrane that is impermeable, gas-permeable, or fluid-permeable can be placed across the circular opening to create a suitable air-liquid interface. This is most often done prior to attaching the plastic ring to the silicone spacer for experimental ease. Once attached, a peristaltic pump or other flow mechanism can be connected to the LSFM-ALI device via tubing through inlet and outlet ports on the side walls of the device. This arrangement allows air exchange when necessary during long-term imaging experiments. Together, this forms a device capable of preserving a pocket of air within a media bath for maintaining samples at an air-liquid interface. The final device design is shown both in fully assembled and exploded views (Figure 1A and B, respectively), as well as the physically realized device itself (Figure 1C). The device is specifically designed for the Multimodal Optical Scope with Adaptive Imaging Correction (MOSAIC)^26^, for which its integration onto the sample holder and into the objective space is also shown (Figure 1D, E). However, the principles of the design can be readily adapted to other upright light sheet systems. The full device design files are available for adaptation (see Data Availability statement).

**Figure 1:**
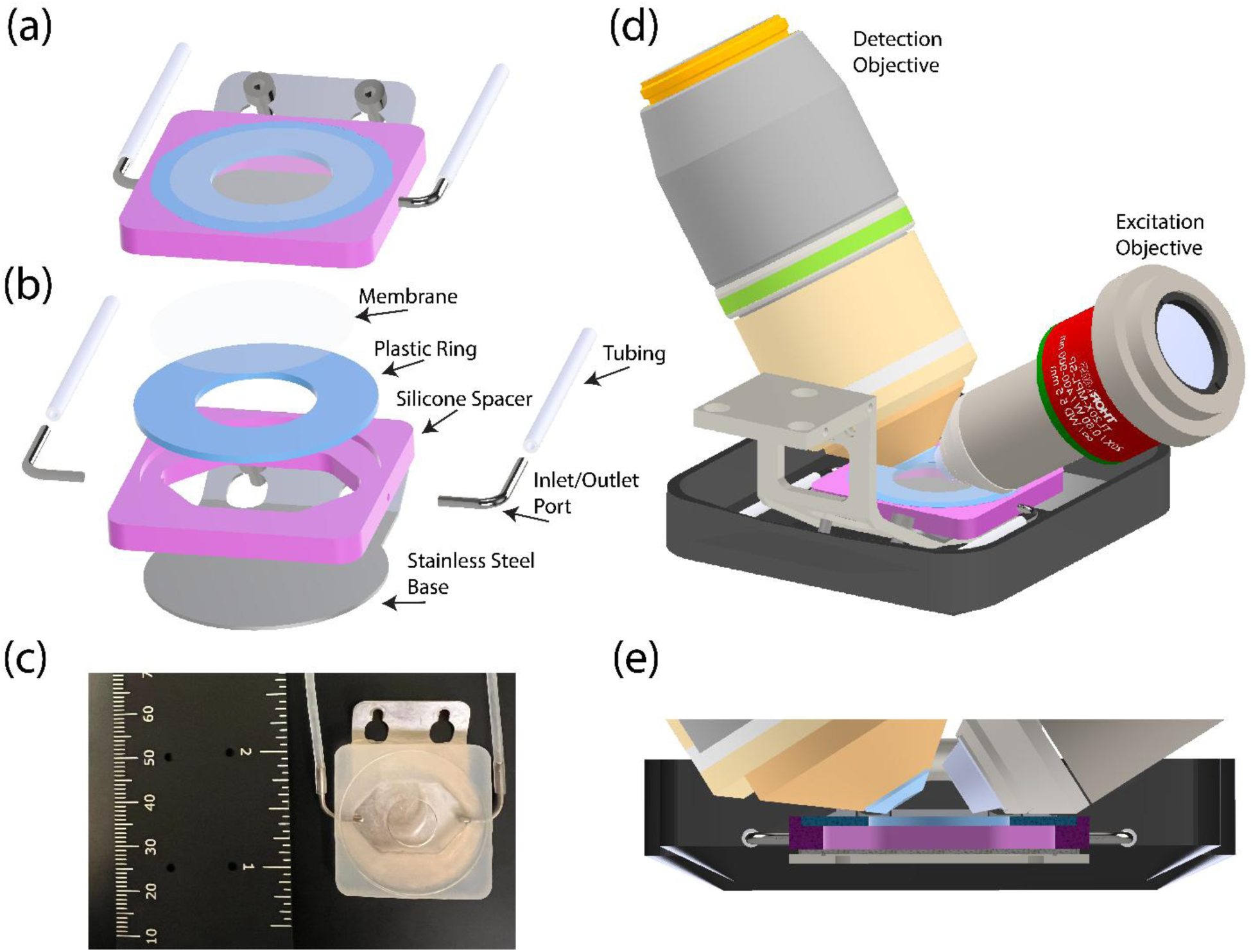
Air-liquid interface (ALI) device for upright light sheet microscopy. (A) Intact and (B) exploded 3D renderings of the LSFM-ALI device. The device consists of a membrane (white/clear), a plastic ring (blue), a silicone spacer (magenta), a stainless-steel base (silver), inlet and outlet ports (silver), and tubing to a peristaltic pump (white). (C) Photo of the LSFM-ALI device with a ruler for scale. (D) 3D rendering showing the LSFM-ALI device together with the sample holder, imaging bucket, and excitation and detection objective lenses. (E) A cross-section view of (D) showing how all components come together at the sample space.

This LSFM-ALI device was subject to several design constraints. First, it needed to fit the current sample holder for the MOSAIC microscope. The sample holder typically secures a 25 mm round coverslip by clamping down a thin, flexible piece of metal via two tightening screws. We leveraged these screws to hold the entire air-liquid interface device as opposed to an intermediate clamp. This setup also made the entire device more easily removable such that samples could be mounted onto the device first and later affixed to the microscope. Next, the device needed to be flexible to accommodate different sample types. To this end, the plastic rings (blue) were machined with multiple inner diameters to allow variations in surface area of the air-liquid interface, and the spacer (pink) was constructed in several different heights to accommodate different volumes of air and sample thicknesses. Third, given the comparatively small volume of air, we found it necessary to include a system to flow air through the device. We therefore incorporated inlet and outlet ports (silver) attached to a peristaltic pump via tubing (white).

### Application 1: *ex vivo* Embryonic Mouse Salivary Glands

A variety of organ explants must be maintained at an air-liquid interface for gas exchange to promote proper growth and development^18–24^. Such an example is an *ex vivo* embryonic mouse salivary gland, generally cultured on floating polycarbonate filters that are not amenable to upright LSFM^18–21^. During development, the epithelium undergoes branching morphogenesis as a means of maximizing functional surface area^18,27^. Here, we used the LSFM-ALI device to enable live LSFM of cell migration during this complex, three-dimensional developmental process.

Embryonic mouse salivary gland explants were mounted onto the device by sandwiching the explants between two gas permeable membranes, which were then affixed to the plastic rings of the LSFM-ALI device (see materials and methods) (Figure 2A, Figure S1). Explants were observed to continue developing in this configuration over multiple days. We first imaged explants from a transgenic mouse line expressing H2B-EGFP to label cell nuclei (Figure 2B, Movie S1). Sustained sample development was unimpeded during 3 hours of imaging, as shown by robust cell divisions (red arrow, Figure 2B, Movie S1). We additionally observed rapid 3D migration of immune cells through the epithelial bud that served to remove cellular debris (Movie S1). LSFM was particularly useful in monitoring these immune cell dynamics given the speed at which they transversed large distances. We next performed multicolor LSFM of explants from WT mice expressing Keratin14-mStayGold and NLS-mScarlet-I3 via lentiviral delivery, marking the intermediate filament cytoskeleton and nuclei, respectively (Figure 2C, Movie S2). During the branching morphogenesis, epithelial cells often traverse narrow intercellular spaces between adjacent epithelial cells (red arrow, Figure 2C, Movie S2). Upright lattice light sheet microscopy enabled us to observe that keratin generally trails the nucleus during migration through these confined regions (magenta, Figure 2C). This demonstration indicates that the LSFM-ALI device can maintain complex specimens for long-term imaging of light sensitive processes.

**Figure 2:**
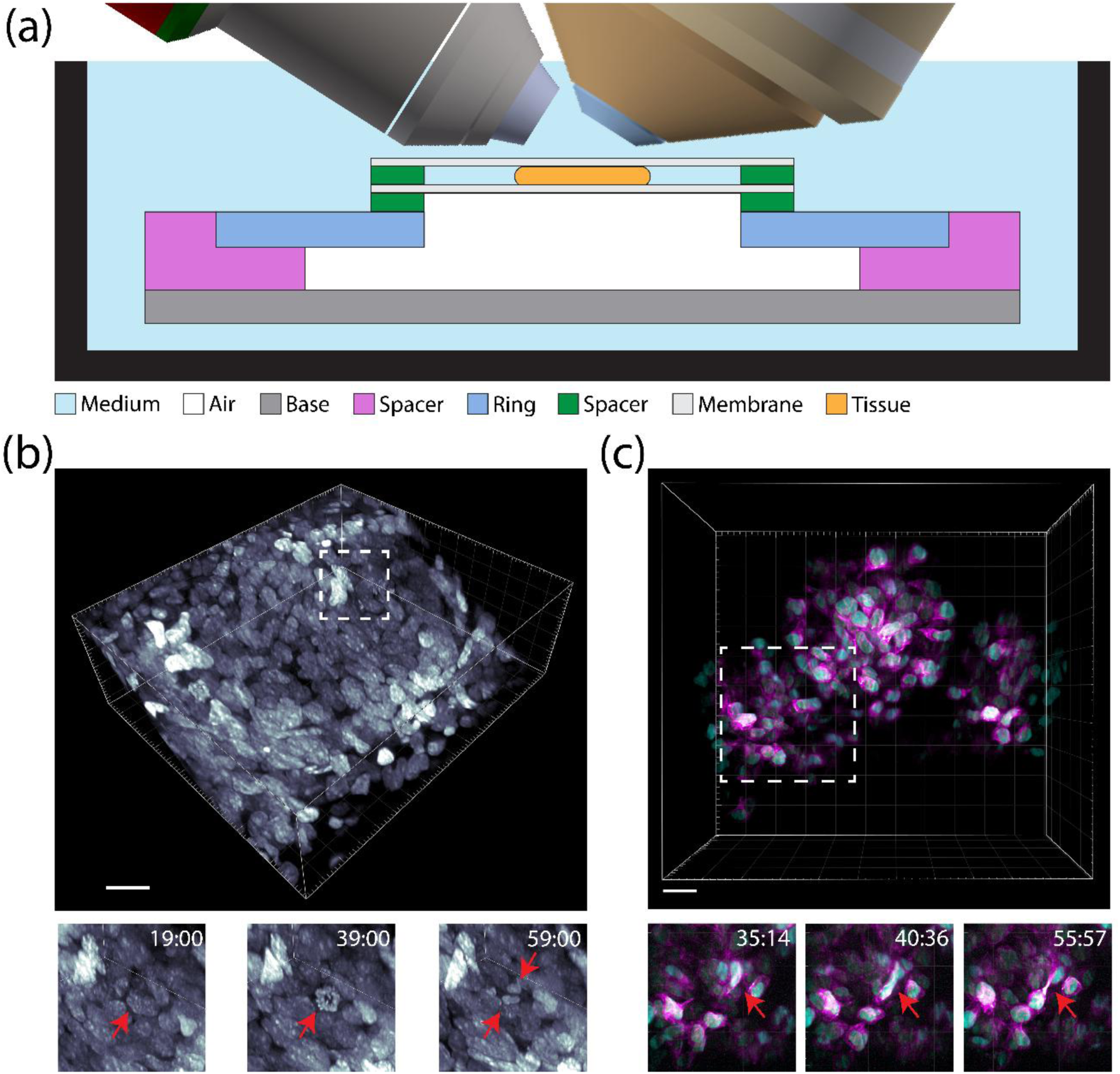
*ex vivo* embryonic mouse salivary gland light sheet microscopy at air-liquid interface. (A) A schematic showing the mounting strategy for the embryonic mouse salivary gland (orange). The gland is submerged entirely in medium with one side exposed to air through a gas-permeable membrane. (B) Volumetric rendering of H2B-EGFP labeled nuclei in the branching epithelium of an embryonic mouse salivary gland (scale bar = 20 μm). Insets show an example of a dividing cell (red arrow). (C) Volumetric rendering of Keratin14-mStayGold (magenta) and NLS-mScarlet-I3 (cyan) in the branching epithelium of an embryonic mouse salivary gland (scale bar = 20 μm). Insets show an example of a migrating cell squeezing through a narrow intercellular space (red arrow).

### Application 2: Human Epidermal Equivalent Culture

Culturing of many epithelial tissues, ranging from airways to skin, requires the basal surface to be exposed to medium while the apical side is exposed to air. This is most commonly achieved by using Transwell membranes^28–30^. The slow remodeling process of these tissues demands sustaining long-term sample viability under imaging conditions. In addition, they pose an exceptional challenge for live, high-resolution imaging due to both tissue complexity and sample mounting. Regarding the latter, two options exist for optically accessing the tissue: apically or basally. When imaging the apical side directly, only an air objective lens may be used, which severely reduces image resolution and further hinders penetration depth due to refractive index mismatch. Such setups also risk the common problem of condensation on the air objective lens during prolonged live imaging experiments. Therefore, it is most practical to image such specimens from the basal side. This is, however, a challenge for upright light sheet configurations.

Our LSFM-ALI device overcomes this barrier. N/TERT-2G keratinocytes^31^ stably expressing ER-StayGold^32^ and mScarlet-I3-K14 were grown at an air-liquid interface on PTFE Transwell membranes to form human epidermal equivalent cultures (see materials and methods) (Figure S2). PTFE was chosen as the membrane material because its refractive index is similar to that of water, and image quality was therefore minimally impacted. Prior to imaging, the Transwell insert itself was cut with a heated scalpel, inverted, and transferred to the plastic ring of the LSFM-ALI device (basal side facing the objective lenses) (Figure 3A, Figure S2). Vacuum grease was used to hold the insert in place and maintain a seal between the insert and the plastic ring. The ring was then moved onto the LSFM-ALI device, which was transferred to the MOSAIC for imaging. We observed cell shape changes through basal and suprabasal layers of the culture, paving the way for future studies on epithelial tissue formation and development (Figure 3B, Movie S3). In addition, we captured the rapid motion of the ER network during extension of the cell body (red arrow, Figure 3C, Movie S4), made possible by the comparatively gentle and high-resolution imaging of upright LSFM.

**Figure 3:**
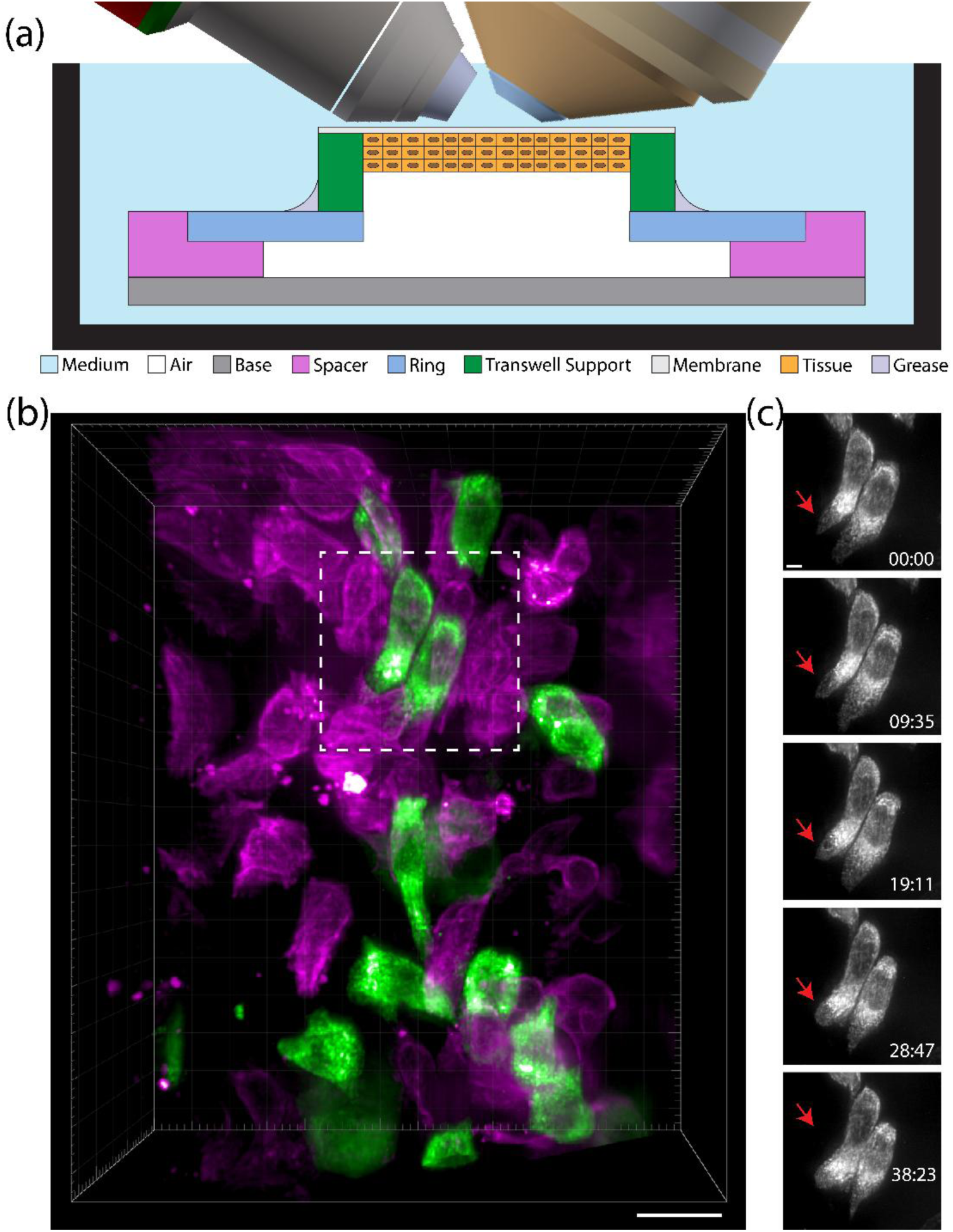
Light sheet fluorescence microscopy of human epidermal equivalent at air-liquid interface. (A) A schematic showing the mounting strategy for the human epidermal equivalent culture. Transwell membranes were cut and affixed to the LSFM-ALI device with vacuum grease such that the culture (orange) was predominately exposed to air with the basal side of the culture exposed to medium through a PTFE Transwell membrane. (B) Volumetric rendering of mScarlet-I3-Keratin14 (magenta) and ER-StayGold (green) labeled cells in a human epidermal equivalent culture (scale bar = 20 μm). (C) Maximum intensity projections of the ER for two cells from the field of view shown in (B) (scale bar = 5 μm). Red arrow indicates a region of cell extension.

### Application 3: in vivo Adult Drosophila melanogaster Brain

Finally, we present a unique use case of imaging *in vivo* adult *Drosophila melanogaster* brains. Live imaging of an adult fruit fly is not compatible with immersion-based systems, thus significantly limiting the available options for rapid, gentle, volumetric optical image acquisition. However, the benefits of upright LSFM could be quite transformative for visualizing light-sensitive processes in the adult brain. The LSFM-ALI device is therefore well suited for addressing these two challenges by providing an environment in which adult flies can be healthily maintained while simultaneously exposing regions of the brain to immersion medium for high-resolution LSFM.

Flies were mounted into aluminum foil sheets pre-attached to the LSFM-ALI plastic rings and the dorsal-posterior cuticle of the head was dissected to expose the brain for imaging (see materials and methods) (Figure S3). The plastic rings were then mounted onto the device, which was submerged in medium on the MOSAIC. This allowed the fly to successfully respirate in a pocket of air while keeping the brain exposed to fluid (Figure 4A). For long term acquisitions (>12 hours), the respiration rate of the fly is sufficient to increase CO2 concentration to a level capable of anesthetization^33^. We therefore maintained constant air flow through the device during experiments. Flies were observed to be alive after experiments by gentling puffing them with air and examining leg movement.

**Figure 4:**
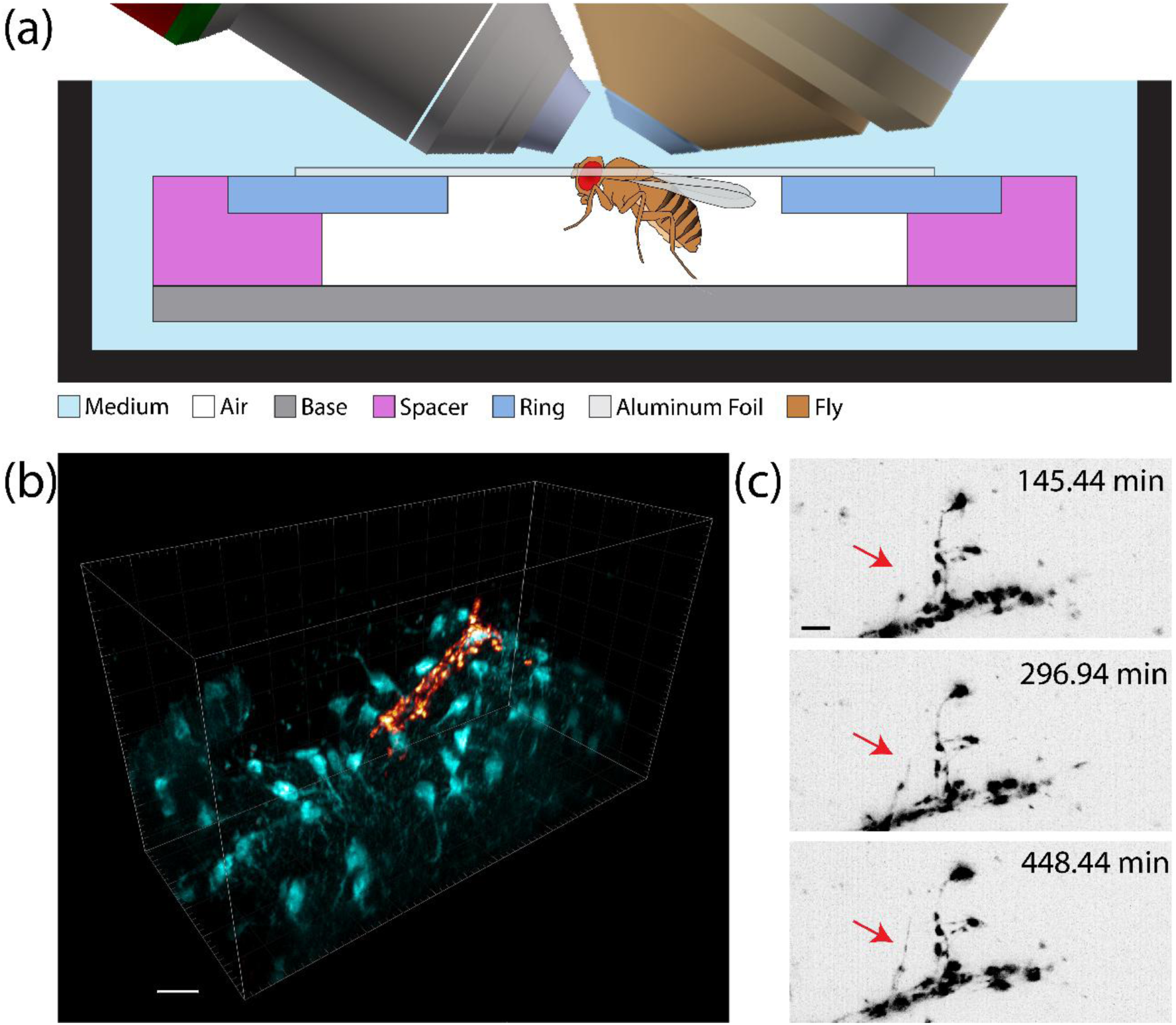
*in vivo* light sheet microscopy of adult *Drosophila melanogaster* brains. (A) A schematic showing the mounting strategy for adult fruit flies. The exposed portion of the brain was kept submerged in medium while the rest of the fly was maintained in an air pocket with fresh airflow to enable respiration. (B) 3D rendering of sLNv dorsal termini labeled with mRFP (orange) and the surrounding astrocytic processes labeled with mIFP (teal) (scale bar = 10 μm). (C) Maximum intensity projection time series of the sLNvs dorsal terminals showing growth of fine processes (red arrow) over the course of several hours (scale bar = 5 μm).

We performed live, multi-channel LLSM of flies expressing mRFP in the small lateral ventral neurons (sLNvs) and mIFP in the astrocyte-like glia. This allowed recording both the sLNv dorsal temini and the surrounding glial processes and cell bodies (Figure 4B, Movie S5). Both neuronal termini from the same brain could be accessed without remounting or additional dissection. sLNvs are known to undergo structural remodeling in a circadian manner, showing a difference in complexity between day and night^34–36^. However, this has been predominantly shown through fixed time point analyses. The device enabled live imaging of part of this process, showcasing the extension of neuronal processes over the course of many hours (red arrow, Figure 4C, Movie S6). Future studies may then examine structural plasticity of the sLNvs, or extend the analysis to other neuronal clusters^37,38^, throughout the circadian cycle within individual animals as opposed to population averaging. This example charts a new path towards imaging non-aquatic animals on upright light sheet microscopes.

## Discussion

Gas exchange across an air-liquid interface is fundamental for many biological processes. Yet, this physical discontinuity can present severe challenges for high-resolution optical imaging. This ultimately limits the range of available tools for understanding an entire class of biological niches. We have presented a device for imaging samples requiring an air-liquid interface on an upright light sheet fluorescence microscope. While the device shown here is optimized for one LSFM implementation, it can serve as a blueprint for designing similar tools for other system designs. With the explicit motivation for flexibility and broad applicability, the LSFM-ALI device is modular in nature, with varied spacer heights and inner ring diameters as well as the integration of fresh air flow through a peristaltic pump system. We demonstrate the utility of this device across three unique biological samples spanning different use cases: *ex vivo* embryonic mouse salivary glands, human epidermal equivalent cultures, and *in vivo* adult *Drosophila* brains. In each instance, we successfully performed volumetric, multi-color, time-lapse LSFM, observing phenomena ranging from cell division and immune cell migration to neuronal process development. These proof-of-concept experiments indicate that more comprehensive imaging-based studies of tissues and animals that were previously not conceivable are now possible. For example, the air flow system could be used to expose adult flies to varied scents while simultaneously monitoring neuronal activity at exceptional speeds or extended durations without light-induced perturbation to the animal.

The LSFM-ALI device, however, is not without limitations. Foremost, sample preparation and mounting are non-trivial endeavors that are often unique for each experiment. Thus, multiple rounds of iteration and testing to ensure sample stability and health are necessary. Additionally, when an experiment requires imaging through a permeable membrane, careful material selection is critical – both for maintaining transport, as well as for its optical properties. In the cases shown here, we found suitable options that minimize their impact on final image quality; however, they still cause slight losses in resolution, light collection efficiency, and working distance. Moreover, while this device enables mounting of potentially large-scale samples, it does not guarantee the entire sample depth can be effectively imaged. Scattering, absorption, and aberrations, which are highly sample-dependent, will ultimately limit light penetration depth to the order of 50-100 μm, regardless of the need for air-liquid interface. Finally, the air perfusion system can cause pulsatile movement of the membrane, which may complicate imaging experiments.

An important point of discussion is the choice of pursuing upright LSFM as opposed to other geometric configurations. We have previously articulated the many advantages of upright LSFM, as well as noted the primary limitation of sample mounting. We believe it is necessary to place this discussion into a broader context. The original light sheet microscope designs are canonically referred to as selective plane illumination microscopes (SPIM)^1^, which feature objectives positioned parallel to the optical table with a specimen held in a vertically orientated tube. This is particularly useful for developmental biology applications. Unfortunately, the geometric constraints make designing a device for imaging a broader spectrum of tissues and organisms at air-liquid interface less tractable. Moreover, these systems are often geared towards lower-resolution, larger-field of view imaging, while our aim here was to preserve higher magnification and resolution. In contrast to the upright configurations, there are also two main classes of inverted LSFM systems: single- or dual-objective. Both are amenable to more standard sample mounting strategies such as multi-well plates or slides, thus improving broad usability. Yet, such dual-objective designs (ex. commercialized LLSM) often require sensitive optics near the specimens and further have comparatively limited working distance. Single-objective designs (ex. oblique plane microscopy (OPM)^39,40^) use multiple tertiary objectives to reimage the specimen. This reduces detection efficiency and results in resolution that varies in all three dimensions. All LSFM configurations are excellent options for high-speed and/or long-term volumetric imaging compared to confocal microscopy, but ultimately the choice of LSFM configuration should be guided by the needs of the experiment^41^. We found the upright LSFM variation to be the most amenable to designing our LSFM-ALI device and thus potentially the most impactful given the broad dissemination of such systems.

Our efforts here are not the first towards extending upright LSFM to a broader application space. Previous works demonstrate microfluidic devices for LLSM^42,43^ that prove useful for imaging weakly adherent cells as well as for precise drug delivery. In addition, several studies have begun examining samples at air-liquid interface using LSFM. Specifically, LSFM through a more classical SPIM geometry has been used to investigate the development of bacterial biofilms^44^, and similar *in vivo* adult *Drosophila* brain imaging has been performed on an upright oblique plane microscope^45^. Our LSFM-ALI device is a complementary approach to these efforts that builds on previous results while also opening new opportunities. This device serves as an important reminder that development of advanced imaging technologies must often be accompanied by innovation at the hardware–biology interface.

## Methods

### LSFM-ALI Device Design

The LSFM-ALI device was custom manufactured by Tokai Hit. The base was machined from SUS304 stainless steel with a thickness of 0.5 mm. The spacer was made from silicone rubber with a hardness of 50 Shore A and was attached to the base via silicone sealant to create a fluid-proof barrier. Spacers were manufactured at 3 different heights of 4 mm, 5 mm, and 6 mm to accommodate multiple sample sizes. Top rings were cut from 1 mm thick polycarbonate sheets. The rings were available with 1 mm, 5 mm, and 10 mm diameter openings to accommodate multiple surface areas, and the outer ring diameter was slightly wider than the inset in the silicone space to ensure a tight seal. Stainless steel pipes of 1.1 mm inner diameter and 1.3 mm outer diameters were used to connect tubing from the peristaltic pump to the LSFM-ALI device. When air flow was not necessary, the side ports were sealed with vacuum grease (Dow Corning, 1597418). The full 3D design files for the LSFM-ALI device are openly available (see Data Availability statement).

### *ex vivo* embryonic Mouse Salivary Gland Preparation

#### Mouse strains

The H2B-EGFP transgenic mouse line FVB(Cg)-Tg(HIST1H2BB-EGFP)1Pa/Jhu was used for nuclear labeling^46^. This line was a gift from A. J. Ewald (Johns Hopkins University), originally obtained from The Jackson Laboratory (RRID:IMSR_JAX:006069) and outcrossed to the FVB/N background. For experiments using wild-type embryos, time-pregnant CD-1 mice were purchased from Charles River.

#### Plasmids

The lentiviral vector was generated by replacing the expression cassette of LentiCRISPR v2 (Addgene, 52961) with a pEF1α short promoter-based construct encoding Keratin14-mStayGold and NLS-mScarlet-I3, separated by a P2A sequence, using Gibson Assembly^47–49^. The resulting plasmid was used for lentivirus production together with packaging plasmids psPAX2 (Addgene, 12260) and pMD2.G (Addgene, 12259).

#### Lentiviral production and titration

Lentivirus was produced, purified, and titrated by the Shared Resource Teams (RRID:SCR_026440) at the Howard Hughes Medical Institute, Janelia Research Campus. Briefly, HEK293T (ATCC, CRL-3216) cells were seeded at 1.5 × 10⁷ cells per 150 cm² dish (10 dishes total) in DMEM (Corning, 17-205-CV) supplemented with 10% fetal bovine serum (FBS; GeminiBio, S11550) and L-glutamine (Thermo Fisher, 25030081). The following day, cells were transfected with the lentiviral plasmid along with psPAX2 (Addgene, 12260) and pMD2.G (Addgene, 12259) packaging plasmids at a ratio of 11.4 μg : 9.1 μg : 2.3 μg per plate using polyethyleneimine (PEI MAX; Kyforabio, 24765-1) in Opti-MEM (Thermo Fisher, 11058021). After 6–8 h, the medium was replaced with DMEM containing 2% FBS.

Viral supernatants were collected at 48 and 72 h post-transfection, pooled, and clarified by low-speed centrifugation followed by filtration through a 0.22 µm filter to remove cellular debris. Virus was concentrated using Centricon 70 centrifugal filters (100 kDa MWCO; MilliporeSigma, UFC710008) and further purified by ultracentrifugation through a 20% sucrose cushion. Viral pellets were resuspended in DMEM/F12 (Thermo Fisher, 11039047), aliquoted, and stored at −80 °C. One aliquot from each preparation was reserved for titration.

For viral titration, 293T cells were plated at 3 × 10^5^ cells per well in a 24-well plate one day prior to transduction. The following day, cells were transduced with virus diluted 1:100 in complete medium containing 8 μg/mL polybrene (MilliporeSigma, TR-1003-G), followed by spinoculation at 1,000 × g for 1 h at 25 °C. Cells were then incubated at 37 °C for at least 2 h before the medium was replaced with fresh complete DMEM, and cultured for an additional 4 days.

On day 4 post-transduction, genomic DNA was isolated using the QIAamp DNA Mini Kit (QIAGEN, 51306). Integration units per mL (IU/mL) were quantified by digital PCR using lentiviral packaging-specific primers/probes and albumin-specific primers/probes on the Absolute Q system (Applied Biosystems, Thermo Fisher). IU/mL was calculated as:

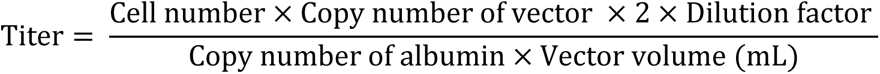

#### Intra-amniotic lentiviral transduction

All surgical procedures for lentiviral delivery were performed by the Shared Resource Teams at the Howard Hughes Medical Institute, Janelia Research Campus. Surgeries were conducted under sterile conditions on timed-pregnant CD-1 dams at embryonic day 10 (E10). Dams were anesthetized with isoflurane (2.5–3.0% in 1 L/min O2 for induction; ∼2% in 0.5–1 L/min for maintenance during surgery). Respiratory rate was monitored throughout the procedure. Buprenorphine (0.1 mg/kg, SC) and ketoprofen (5 mg/kg, SC) were administered preoperatively, and ophthalmic lubricant was applied. Animals were positioned supine on a heated pad set to 41 °C, and the lower abdomen was shaved and disinfected.

A sterile drape with a central opening was placed over the surgical site, and warm saline was applied to the exposed region. A midline laparotomy was performed, followed by incision of the linea alba to access the peritoneal cavity. The uterine horns were gently exteriorized onto the drape. Embryos were stabilized using ring and Adson forceps. Glass micropipettes with a tip diameter of 30–40 μm were backfilled with mineral oil, mounted onto a Nanoject III injector (Drummond Scientific), and front-loaded with lentiviral suspension mixed with Fast Green. Prior to injection, the micropipette tip was moistened with saline to ensure patency. The micropipette was inserted through the uterine wall into the amniotic cavity under visual guidance, and 1–5 μL of lentiviral suspension (2.9 × 10^10^ IU/mL) was delivered using preprogrammed pulses on the Nanoject III. After injections were completed for all embryos, the uterine horns were returned to the abdominal cavity.

The abdominal wall was closed with interrupted 5-0 absorbable sutures, and the skin was closed using a continuous intradermal pattern. Marcaine (<7 mg/kg) was administered subcutaneously along the incision prior to completing closure. Vetbond was applied to seal the incision. Isoflurane was discontinued at the end of surgery, and animals were maintained on oxygen and heat support until fully ambulatory. Postoperative ketoprofen (5 mg/kg, SC) was administered once daily for two days. Dams were monitored for at least three days for signs of distress, bleeding, or wound dehiscence.

#### Salivary gland isolation and culture

Submandibular salivary glands were dissected from E13 embryos derived from either the H2B-EGFP transgenic line or CD-1 wild-type embryos subjected to intra-amniotic lentiviral injections, as previously described^50^. In brief, mouse embryos were decapitated using a scalpel (Fine Science Tools, 10011-00 and 10003-12). The detached head was held laterally by inserting one prong of forceps (Fine Science Tools, 11251-20) through the dorsal surface, and the scalpel was used to cut across the oral opening to separate the mandible and tongue region, where the submandibular glands are located.

Under a dissecting scope, the isolated mandible was transferred to a glass surface with the tongue oriented downward. The mandible was bisected along the midline using forceps to expose the tongue and the paired submandibular glands attached at its base. Surrounding tissues were carefully removed, and the glands were gently detached with forceps and transferred into a 35-mm dish containing 2 mL of DMEM/F12 (Thermo Fisher, 11039047) supplemented with 150 μg/mL vitamin C (MilliporeSigma, A7506), 50 μg/mL transferrin (MilliporeSigma, T8158) and 1x PenStrep (100 units/mL penicillin, 100 μg/mL streptomycin; Thermo Fisher, 15140163).

#### Sample mounting

To mount submandibular salivary glands for air-liquid interface culture (Figure S1), a two-layer lumox sandwich assembly was constructed. To prepare the base layer, an 18 mm × 18 mm double-sided spacer with a 9 mm opening diameter (Grace BioLabs, 654002) was adhered to the underside of the lumox film of a 35-mm Sarstedt dish (Sarstedt, 94.6077.333) (Figure S1A). After attachment, the lumox film together with the spacer was excised from the dish using a scalpel and mounted onto the plastic ring of the LSFM-ALI device, with a 10 mm diameter central aperture, forming the base layer of the assembly (Figure S1B). For the upper layer, two additional rectangular spacers (∼ 5 mm × 5 mm) were cut and attached to two opposing sides of the bottom surface of a second lumox film (Figure S1C). This lumox film was then trimmed into an hourglass shape, with the narrowed region positioned between the two spacer squares (Figure S1D). The geometry of this top layer provides gentle mechanical confinement of the gland while preserving access to culture medium.

The hourglass-shaped lumox film was adhered to the base lumox layer, forming a sandwich assembly with a 120 μm interlayer gap defined by the spacer thickness (Figure S1E). Approximately 3 μL of complete DMEM/F12 medium was placed next to the narrowed region of the hourglass-shaped lumox film. Salivary glands were transferred into this droplet and carefully positioned between the lumox layers using forceps (Figure S1F,G). An additional 100 μL of medium was added on top of the upper lumox layer to prevent drying. The complete assembly was placed onto the LSFM-ALI device with the base lumox layer facing the air chamber, and the hourglass-shaped lumox layer facing upward toward the culture medium (Figure S1H). After mounting onto the microscope, 40 mL of culture medium was filled in the imaging chamber. Because lumox is gas permeable, this configuration allowed oxygen diffusion from the air chamber below while preventing medium from entering the compartment.

### Human Epidermal Equivalent Culture Preparation

#### Lentivirus generation

The ER-StayGold (er-(n2)oxStayGold(c4)v2.0)^51^ and mScarlet-I3-K14 sequences were inserted into the pLV lentiviral vector backbone and synthesized by VectorBuilder. The sequences of all plasmids were obtained from VectorBuilder. Whole Plasmid Sequencing was performed by Plasmidsaurus using Oxford Nanopore Technology with custom analysis and annotation to confirm sequences. Lentiviruses were made by co-transfection into human embryonic kidney-293FT cells with pMD2.G (encoding VSV-G) and psPAX2 (encoding Gag and Pol) and collection of culture supernatants 24‒72 h after transfection. Lentivirus was concentrated with the Lenti-X Concentrator kit (Takara Bio, 631231) following the manufacturer’s protocol. pMD2.G and psPAX2 (gifts from Didier Trono, Addgene, 12259; Addgene, 12260, respectively).

#### Cell line generation

N/TERT-2G keratinocytes^31^ were cultured in Keratinocyte Serum-Free Media (K-SFM) supplemented with 30 μg/mL bovine pituitary extract and 0.2 ng/mL human recombinant epidermal growth factor (Gibco, 37010022,). N/TERTs were transduced with lentivirus by incubating cells with 8–10 μg/mL polybrene (EMD Millipore, TR-1003-G) in K-SFM medium for 24–48 h. Bulk sorting of cell lines expressing the various constructs was performed by fluorescence-activated cell sorting to obtain populations with optimal expression levels, as determined by localization of ER-StayGold and mScarlet-I3-K14.

#### Preparation of Transwell inserts

Transwell inserts (Corning, CLS3495) were aseptically placed in the provided 24-well plates. 100 µl of K-SFM media was added to the top of the inserts while 600 µl of K-SFM media was added to the bottom. The 24-well plate was then placed in a cell culture incubator (at 37⁰C and 5% CO2) for an hour.

#### Subculturing

For passaging, cells were quickly washed once with 1X PBS- before incubating at 37°C and 5% CO2 with fresh 1X PBS- for 10 minutes. The PBS- was then replaced with 2 ml of TrypLE Express (Gibco, 12604-013), and cells were incubated at 37⁰C and 5% CO2 for 7 minutes. Following visual confirmation of dissociation, cells were centrifuged at 0.2rcf for 5 minutes. PBS was aspirated, and cells were resuspended in K-SFM media, followed by counting and seeding for experiments.

#### 3D cell culture

Medium on top of Transwell inserts was aspirated. After manual counting with a hemocytometer, 100,000 cells were seeded in each insert. Cells reached 100% confluency after two days. 2D growth medium (K-SFM) above and below the insert was replaced with 3D growth medium, i.e., CNT-Prime Epithelial 3D Airlift Medium (CELLnTEC, CnT-PR-3D). Following the overnight acclimation period, the medium above the insert was aspirated without replacement. The 3D growth medium below the inserts was refreshed daily until cells were imaged after 5 days of air-liquid interface culture.

#### Sample mounting

To maintain structural integrity of the fragile PTFE membrane, a flame-heated #10/#11 scalpel was used to cut the polystyrene support above the PTFE membrane (Figure S2A-C). A thin layer of vacuum grease (Dow Corning, 1597418) was applied to the top side of the plastic ring of the LSFM-ALI device (Figure S2D). The cut Transwell membrane was inverted and attached to a plastic ring via the vacuum grease, creating a water-tight seal (Figure S2E). The plastic ring was then loaded onto the rest of the LSFM-ALI device (Figure S2F), which was then mounted onto the MOSAIC system. Imaging was performed in CNT-Prime Epithelial 3D Airlift Medium with additional antibiotics/antimycotics (Sigma, P0781) at 37⁰C with 5% CO2. After the experiment, it was confirmed that no medium had entered the air pocket of the LSFM-ALI device.

### Adult Drosophila melanogaster Preparation

#### Fly stocks

All flies were raised and maintained at 25°C in vials containing standard cornmeal yeast agar medium under 12:12 h light:dark (LD) cycles. All experiments were performed with 3-5 days old adult flies randomly picked from their vials. Stocks white1118 (#5905), ;;GMR86E01-Gal4 (#45914) and ;;UAS-mIFP (#64181) were obtained from Bloomington Stock Center. *Pdf*-RFP was generously provided by Justin Blau^52^.

#### Sample mounting and dissection

Prior to dissection, the central holes in the plastic rings for the LSFM-ALI device were sealed aluminum foil (Figure S3A). Pyramid-shaped holes were then cut into the foil to provide locations for the flies to be mounted (Figure S3A). Each fly was briefly anesthetized in an empty vial on ice and its head and thorax were gently positioned into the pyramid-shaped hole in the aluminum foil using forceps under a dissection microscope (Figure S3B). The plastic ring was then attached to the rest of the LSFM-ALI device. The fly head was bent forward to give access to its posterior surface (Figure S3C,D), and it was then carefully immobilized by the eyes using beeswax (70°C; Sigma, 243248) and a thin-end wax carving pencil (Renfert, 20348) (Figure S3E). The thorax was pushed down to ensure that the spiracles remained within the air chamber of the LSFM-ALI device and subsequently immobilized by its posterior end. For short-term imaging (< 3 h), the dorsal side of the foil was immersed in dissection medium (103 mM NaCl, 3 mM KCl, 5 mM TES, 8 mM trehalose, 10 mM glucose, 26 mM NaHCO3, 1 mM NaH2PO4, 4 mM MgCl2 and 1.5 mM CaCl2), while the ventral side was kept dry. An incision was made in the dorso-posterior region of the head using a tungsten needle (Fine Science Tools). The cuticle and the air sacs were removed using superfine forceps (Fine Science Tools, 11252-00), thereby enabling optical access to the dorsal protocerebrum (Figure S3F). The M16 muscles were meticulously sectioned to mitigate brain movement. For long-term imaging (> 12 h), dry dissection was quickly performed, and the window was sealed by spreading Kwik-Sil silicone (World Precision Instruments) along the head with a tungsten needle. In both cases, imaging was carried out in the dissection medium as the immersion medium for the objectives at ambient environmental conditions. Air flow was maintained during long-term imaging experiments via a peristaltic pump (Tokai Hit).

### Imaging

All imaging was performed on the MOSAIC system^26^ housed at HHMI Janelia Research Campus’ Advanced Imaging Center using the Lattice Light Sheet Microscopy (LLSM) imaging modality^10^. Excitation was performed using a square lattice pattern and a Thorlabs TL20x-MPL, 0.6 NA objective lens. Image stacks were acquired by scanning the sample stage either horizontally at an angle of 32.45° relative to the optical axis of the detection objective (X stage scanning) or directly in line with the optical axis of the detection objective (XZ stage scanning). Emitted fluorescence collected through a detection objective (Zeiss Plan-Apo 20x, NA 1.0 DIC M27 75 mm) and was directed to two detectors (Hamamatsu ORCA-Fusion) via a Semrock Di03-R561-t3-32 × 40 dichroic mirror. Subsequently, light reflected from the dichroic mirror was filtered through a green bandpass filter (Semrock, FF01-520/35-25), and light transmitted through the dichroic was filtered through notch filters (Semrock, NF03-561E-25; Semroc, NF03-642E-25). As previously described^26,53^, a baseline correction for aberrations through microscope optics was applied to the deformable mirror (ALPAO, DM-69) for all imaging experiments. For specific experiments, subsequent adaptive optics corrections were applied as previously described^26,53^. In brief, a femtosecond laser (Coherent Chameleon LS II, tuned to 920 nm) was focused through the detection objective and scanned through the desired image volume. The emitted fluorescence was passed through a custom Shack-Hartmann wavefront sensor to measure the aberration to the wavefront, after which the deformable mirror was updated to correct for this aberration. This process was repeated for three iterations prior to imaging. Full details for each imaging experiment are provided in Supplemental Table 1.

### Image Processing

Each dataset required varied processing approaches, which are explicitly detailed in Supplementary Table 1. For data collected using X stage scanning, deskewing is necessary to restore the image stack to its correct geometric alignment. In some instances, sample motion required alignment between adjacent frames in a given volume (intra-stack registration). This was accomplished using the phase cross correlation registration function in scikit-image between each slice in the stack. Registration was performed on a reference channel and then applied to the other channel, as described at https://github.com/aicjanelia/visitor-ceriani. Once in the proper alignment, all data sets were deconvolved with 10 iterations of Richardson-Lucy deconvolution using experimentally collected point spread functions for each excitation wavelength. Deskewing and deconvolution were performed on the compute cluster at HHMI Janelia Research Campus using custom software^54^. For visualization, 3D renderings were generated in Imaris (Oxford Instruments), and 2D representations were generated in FIJI^55^. Any non-linear intensity modifications or drift corrections for visualization purposes are noted in Supplementary Table 1.

## Data Availability

The 3D design file for the LSFM-ALI device as well as all raw image files underlying the images and movies shown here are available in the following repository: https://doi.org/10.25378/janelia.c.8403681.

## Code Availability

The software for intrastack registration is available at https://github.com/aicjanelia/visitor-ceriani. The software for deskewing and deconvolution is available at https://github.com/aicjanelia/LLSM.

## Acknowledgements

We thank: the Molecular Genomics and Viral Tools support teams at Janelia for plasmid prep and lentivirus packaging; Janelia Experimental Technology for fabrication assistance; Janelia Invertebrate Shared Resource for maintaining fly stocks; the Jayaraman and Shroff labs at Janelia Research Campus for thoughtful discussion and equipment loans, the full Janelia Advanced Imaging Center, Integrative Imaging, and Light Microscopy teams for thoughtful discussion on this project; Dan Milkie for assistance with microscope control software; J. Bednarczyk and staff from Penn State College of Medicine’s Flow Cytometry Core for assistance with cell sorting.

This work was supported by: Howard Hughes Medical Institute; Children’s Miracle Network Faculty Research grant 1110000016 (NKB); NIH grant R01AR081883 (APK); NIH grant R01AR048266 (APK); NIH MIRA grant R35GM150921 (SNS); graduate fellowships from the Argentine Research Council for Science and Technology (CONICET) (JII, MRC, and FJT; MFC is a member of CONICET); Agencia I+D+i grant PICT2018-0995 (MFC), NIH grant R01NS108934 (MFC).

## Author Contributions

CMH, JA, TLC, KI, and TT designed the LSFM-ALI device. KI and TT fabricated the LSFM-ALI device. CMH, EQ, EAM, and SW performed the *ex vivo* mouse embryonic salivary gland experiments. SY developed the Keratin14-mStayGold and NLS-mScarlet-I3 construct. CMH and WG performed the human epidermal equivalent culture experiments. NKB assisted in generating the necessary cell lines for the human epidermal equivalent cultures. CMH, JII, MRC, and FT performed the *in vivo* adult *Drosophila melanogaster* brain experiments. CMH, RML, and OFP processed data sets. MFC, APK, SNS, SW, and TLC contributed funding and resources. All authors contributed to the writing and editing of the manuscript.

