## Supplemental Materials for "Adapting Upright Light Sheet Fluorescence Microscopy for Imaging at Air-Liquid Interface"

#### Supplemental Figures

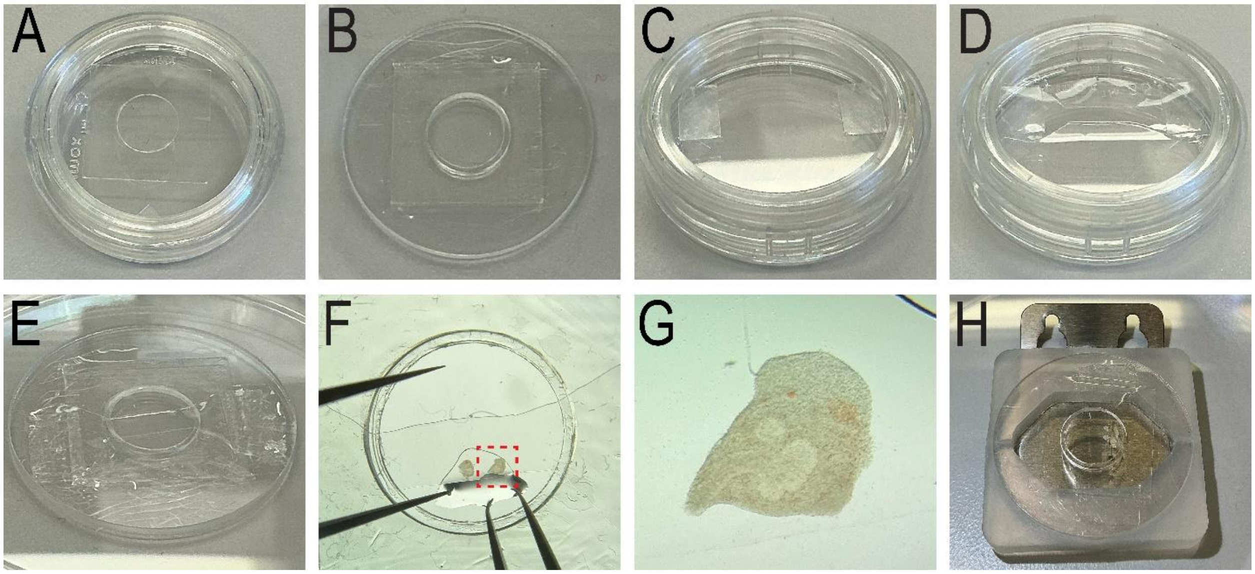


Figure S1: Mounting of embryonic mouse salivary glands. (A) A spacer is attached to the underside of a lumox film. (B) The film and spacer are removed and attached to the LSFM-ALI device's plastic ring. (C) Rectangular spacers are attached to the underside of a second lumox film. (D) The film was trimmed into an hourglass shape with the narrow region between the two spacers. (E) The hourglass-shaped assembly is adhered to the first lummox film assembly. (F) Embryonic mouse salivary gland explants are positioned in a drop of medium under the hourglass lumox film to provide gentle mechanical confinement. (G) A zoom in of the red dashed gland from (F). (H) The plastic ring is mounted onto the rest of the LSFM-ALI device.

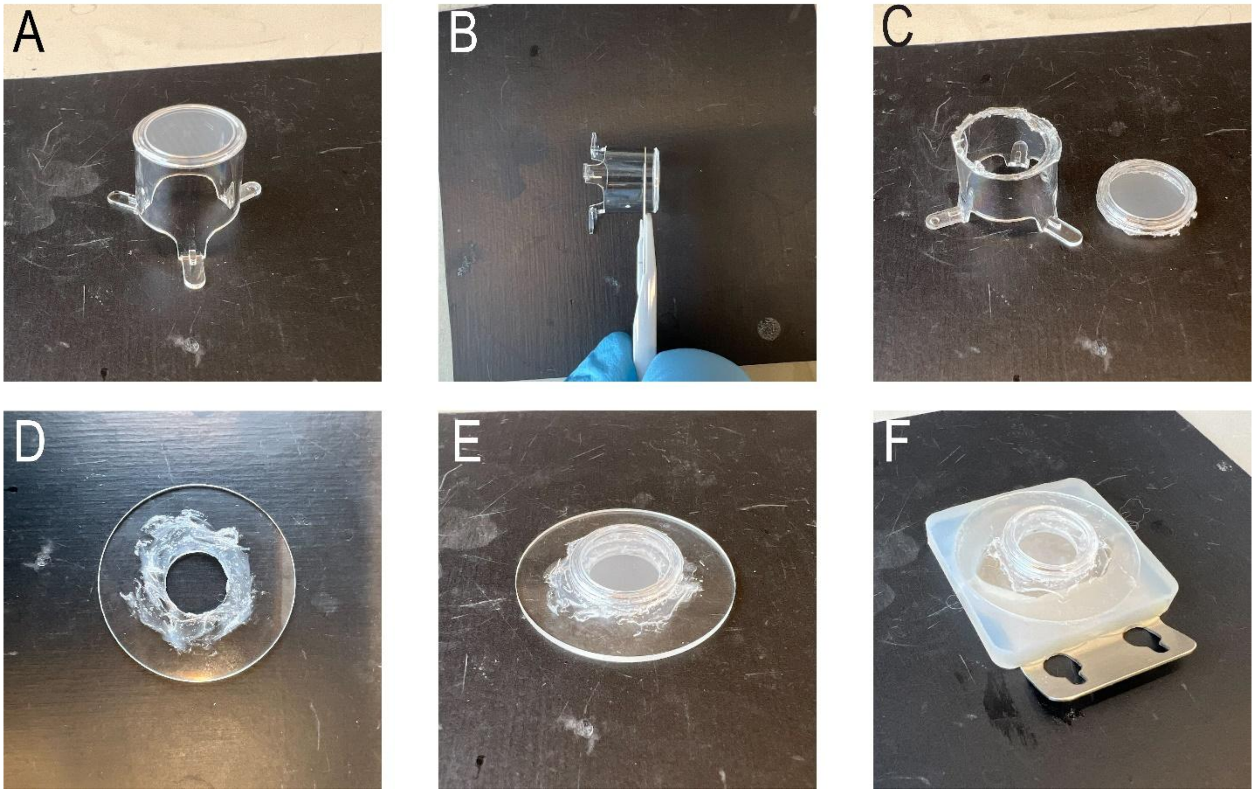


Figure S2: Mounting of human epidermal equivalent cultures. (A) A Transwell membrane insert is removed from the multi well plate. (B) A heated scalpel is used to cut the plastic below the membrane. (C) The tension on the membrane is kept intact as it is removed from the rest of the support. (D) A ring of vacuum grease is applied to the plastic ring of the LSFM-ALI device. (E) The membrane is attached to the greased ring to create a watertight seal. (F) The plastic ring is attached to the rest of the LSFM-ALI device. Note that shown here is the procedure performed on a blank Transwell membrane, mounting of actual samples was performed aseptically in a hood.

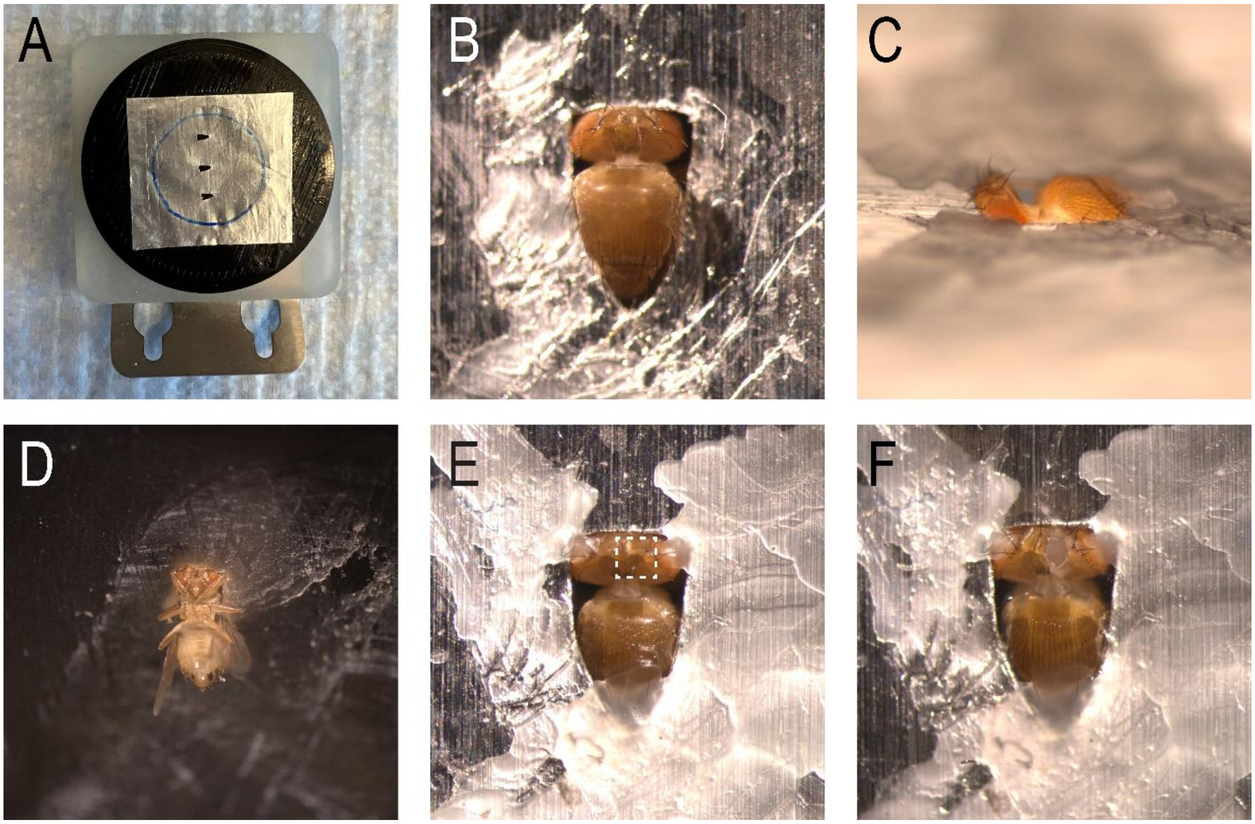


Figure S3: Mounting of adult *Drosophila melanogaster*. (A) Aluminum foil is pre-attached to the LSFM-ALI device plastic ring. Pyramid-shaped holes are cut into the foil. (B) Flies are mounted through the pyramid-shaped opening. Side view (C) and underside view (D) show the positioning of the fly in the hole. (E) Beeswax is used to keep the fly in place. (F) The white dashed region in (E) is dissected to expose the brain for imaging.

#### Supplemental Movie Captions

Movie S1: *ex vivo* embryonic mouse salivary gland light sheet microscopy at air-liquid interface. Volumetric rendering of H2B-EGFP labeled nuclei in the branching epithelium of an embryonic mouse salivary gland (scale bar = 20 μm).

Movie S2: *ex vivo* embryonic mouse salivary gland light sheet microscopy at air-liquid interface. Volumetric rendering of Keratin14-mStayGold (magenta) and NLS-mScarlet-I3 (cyan) in the branching epithelium of an embryonic mouse salivary gland (scale bar = 20 μm).

Movie S3: Light sheet fluorescence microscopy of human epidermal equivalent at air-liquid interface. Volumetric rendering of mScarlet-I3-Keratin14 (magenta) and ER-StayGold (green) labeled cells in a human epidermal equivalent culture (scale bar = 20 μm).

Movie S4: Light sheet fluorescence microscopy of human epidermal equivalent at air-liquid interface. Maximum intensity projections of the ER-StayGold for two cells from broader field of view shown in Movie S3 (scale bar = 5 μm).

Movie S5: *in vivo* light sheet microscopy of adult *Drosophila melanogaster* brains. 3D rendering of sLNv dorsal termini labeled with mRFP (orange) and the surrounding astrocytic processes labeled with mIFP (teal) (scale bar = 10 μm).

Movie S6: *in vivo* light sheet microscopy of adult *Drosophila melanogaster* brains. Maximum intensity projection time series of the sLNvs dorsal terminals showing growth of fine processes over the course of several hours (scale bar = 5 μm).

### Supplemental Tables

Table S1: Sample details, imaging parameters, and processing steps for all experiments.

| Figure | Sample | Environmental Conditions | Fluorescent Labels | Channels | Scan Type | Raw Voxel Size (µm) | Raw Field of view (voxels) | Exposure Time (ms) | Time Points | Temporal Sampling | Lattice Pattern | AO? | Processing | Visualization |
|---|---|---|---|---|---|---|---|---|---|---|---|---|---|---|
| Figure 2B, Movie 1 | Mouse embryonic salivary gland explant | 37°C; 5% CO2 | H2B-EGFP | 488 nm | X Stage Scanning | .108 x .108 x .5 | 512 x 1500 x 201 | 20 | 300 | 60.0 s | Square; NA Max: 0.4; NA Min: 0.34; Cropping: 10; Envelope: 4 | Yes | Deskew; Deconvolution: 10 iterations of Richardson-Lucy with experimental PSF | Imaris |
| Figure 2C, Movie 2 | Mouse embryonic salivary gland explant | 37°C; 5% CO2 | NLS-mScarlet3; keratin-mStayGold | 488 nm; 560 nm | X Stage Scanning | .108 x .108 x .5 | 512 x 2048 x 351 | 20 | 125 | 62.2 s | Square; NA Max: 0.4; NA Min: 0.34; Cropping: 10; Envelope: 10 | Yes | Deskew; Deconvolution: 10 iterations of Richardson-Lucy with experimental PSF | Imaris |
| Figure 3B,C, Movie 3,4 | Human epidermal equivalent culture (N/TERT keratinocytes) | 37°C; 5% CO2 | mScarlet-I3-keratin14; ER-StayGold | 488 nm; 560 nm | X Stage Scanning | .108 x .108 x .5 | 512 x 1536 x 201 | 10 | 123 | 23.0 s | Square; NA Max: 0.4; NA Min: 0.34; Cropping: 10; Envelope: 10 | Yes | Deskew; Deconvolution: 10 iterations of Richardson-Lucy with experimental PSF; Figure 3B and Movie 3 - Gamma = 1.5 (Imaris); Figure 3C and Movie 4 - Gamma = 0.5 (FJI) | Imaris, FIJI |
| Figure 4B, Movie 5 | Adult *Drosophila melanogaster* | Ambient environment | *Pdf*-RFP; UAS-mIFP | 560 nm; 642 nm | XZ Stage Scanning | .108 x .108 x .25 | 512 x 1536 x 301 | 20 | 1 | N/A | Square; NA Max: 0.4; NA Min: 0.34; Cropping: 10; Envelope: 8 | No | Intra-stack registration; Deconvolution: 10 iterations of Richardson-Lucy with experimental PSF | Imaris |
| Figure 4C, Movie 6 | Adult *Drosophila melanogaster* | Ambient environment | *Pdf*-RFP; UAS-mIFP | 560 nm; 642 nm | XZ Stage Scanning | .108 x .108 x .25 | 512 x 1536 x 301 | 20 | 101 | 363.5 | Square; NA Max: 0.4; NA Min: 0.34; Cropping: 10; Envelope: 4 | No | Intra-stack registration; Deconvolution: 10 iterations of Richardson-Lucy with experimental PSF; 3D drift correction (FIJI) | FIJI |
